# Directed robust generation of functional retinal ganglion cells from Müller glia

**DOI:** 10.1101/735357

**Authors:** Dongchang Xiao, Suo Qiu, Xiuting Huang, Rong Zhang, Qiannan Lei, Wanjing Huang, Haiqiao Chen, Bin Gou, Xiaoxiu Tie, Sheng Liu, Yizhi Liu, Kangxin Jin, Mengqing Xiang

## Abstract

Glaucoma and optic neuropathies cause progressive and irreversible degeneration of retinal ganglion cells (RGCs) and the optic nerve and are currently without any effective treatment. Previous research into cell replacement therapy of these neurodegenerative diseases has been stalled due to the limited capability for grafted RGCs to integrate into the retina and project properly along the long visual pathway to reach their brain targets. In vivo RGC regeneration would be a promising alternative approach but mammalian retinas lack regenerative capacity even though cold-blood vertebrates such as zebrafish have the full capacity to regenerate a damaged retina using Müller glia (MG) as retinal stem cells. Nevertheless, mammalian MG undergo limited neurogenesis when stimulated by retinal injury. Therefore, a fundamental question that remains to be answered is whether MG can be induced to efficiently regenerate functional RGCs for vision restoration in mammals. Here we show that without stimulating proliferation, the transcription factor (TF) Math5 together with a Brn3 TF family member are able to reprogram mature mouse MG into RGCs with exceedingly high efficiency while either alone has no or limited capacity. The reprogrammed RGCs extend long axons that make appropriate intra-retinal and extra-retinal projections through the entire visual pathway including the optic nerve, optic chiasm and optic tract to innervate both image-forming and non-image-forming brain targets. They exhibit typical neuronal electrophysiological properties and improve visual responses in two glaucoma mouse models: *Brn3b* null mutant mice and mice with the optic nerve crushed (ONC). Together, our data provide evidence that mammalian MG can be reprogrammed by defined TFs to achieve robust in vivo regeneration of functional RGCs as well as a promising new therapeutic approach to restore vision to patients with glaucoma and other optic neuropathies.

---

Glaucoma is a neurodegenerative disorder characterized by progressive and irreversible degeneration of RGCs and the optic nerve, and is the second leading cause of blindness worldwide(*1*, *2*). Despite its discovery almost a century and half ago(*3*), there is currently still no effective treatment for glaucoma. In vivo RGC regeneration would be an ideal therapy but mammalian retinas are thought to lack regenerative capacity. In spite of this, the MG have been shown unambiguously to serve as retinal stem cells to repair injured retinas in zebrafish(*4*, *5*). Similarly, mammalian MG display stem cell-like/late retinal progenitor features, e.g. having a molecular signature similar to late retinal progenitors(*6*, *7*), exhibiting limited proliferative and neurogenic capacity in damaged retinas(*8*–*12*), and transdifferentiating into rods by a combination of b-catenin and TFs(*13*). We sought to harness the stem cell-like property of MG to regenerate RGCs in vivo by TF-directed reprogramming. During murine retinogenesis, the bHLH TF Math5/Atoh7 is transiently expressed in a subset of retinal progenitors and required for conferring them with the competence of RGC generation(*14*–*16*) (Fig. 1A). Previously, we and others have demonstrated the expression of the POU-domain transcription factor Brn3b/Pou4f2 in RGC precursors and its crucial function in RGC specification and differentiation(*16*–*21*) (Fig. 1A). We thus investigated the ability of the Math5 and Brn3b TF combination to reprogram adult mouse MG into RGCs.

**Fig. 1.**
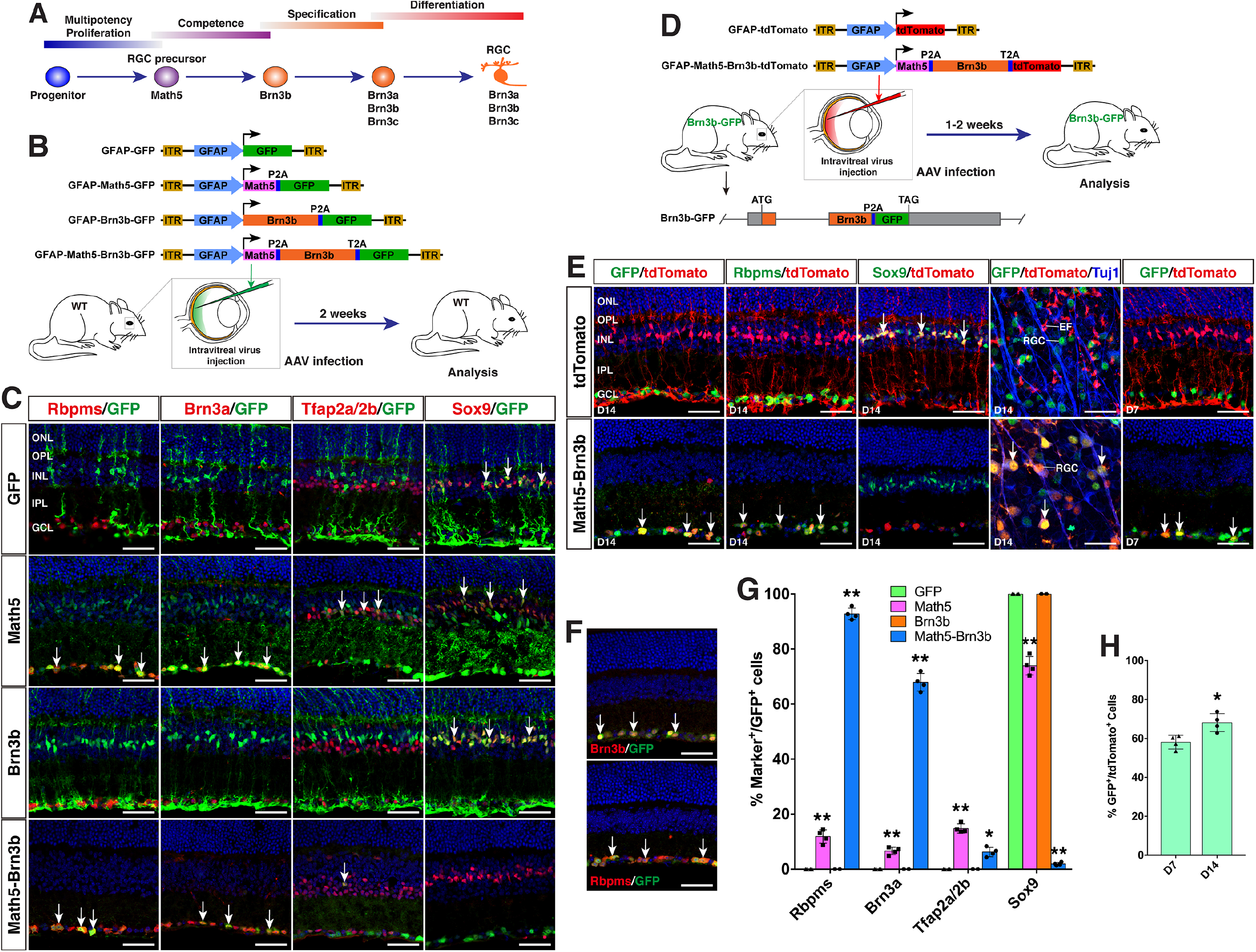
Generation of RGCs by TF-mediated reprogramming of adult mouse MG. (**A**) Schematic illustration of developmental stages and the associated Math5 and Brn3 TFs leading to RGC specification and generation from retinal progenitors. (**B**) Schematic of the AAV constructs and infection procedure to generate RGCs in adult wild-type (WT) mice. (**C**) Two weeks after intravitreal injection of GFAP-GFP, GFAP-Math5-GFP, GFAP-Brn3b-GFP or GFAP-Math5-Brn3b-GFP AAVs, sections from infected retinas were double-immunolabeled with the indicated antibodies and counterstained with nuclear DAPI. Arrows point to representative colabeled cells. (**D**) Schematic of the AAV constructs and infection procedure to generate RGCs in Brn3b-GFP reporter mice. (**E**) One (D7) or two (D14) weeks after intravitreal injection of GFAP-tdTomato or GFAP-Math5-Brn3b-tdTomato AAVs, sections or flat-mounts from infected retinas were double-immunolabeled with the indicated antibodies. Sections were also counterstained with nuclear DAPI. Note that fluorescent staining for Rbpms and Sox9 is in manually assigned false green color. Arrows point to representative colabeled cells. (**F**) Sections from retinas of Brn3b-GFP reporter mice without AAV infection were double-immunolabeled with the indicated antibodies and counterstained with nuclear DAPI. Arrows point to representative colabeled cells. (**G**) Quantitation of GFP+ cells that become immunoreactive for Rbpms, Brn3a, Tfap2a/2b or Sox9 in wild-type retinas infected with GFAP-GFP, GFAP-Math5-GFP, GFAP-Brn3b-GFP or GFAP-Math5-Brn3b-GFP AAVs. Data are presented as mean ± SD (n=4). Asterisks indicate significance in unpaired two-tailed Student’s t-test: *p<0.0005,**p<0.0001. (**H**) Quantitation of tdTomato+ cells that become immunoreactive for GFP in Brn3b-GFP reporter retinas one or two weeks following infection with GFAP-tdTomato or GFAP-Math5-Brn3b-tdTomato AAVs. Data are presented as mean ± SD (n=4). Asterisks indicate significance in unpaired two-tailed Student’s t-test: *p<0.05. Abbreviations: EF, MG endfoot; GCL, ganglion cell layer; INL, inner nuclear layer; IPL, inner plexiform layer; ONL, outer nuclear layer; OPL, outer plexiform layer; RGC, retinal ganglion cell. Scale bars: 40 μm (C, E, F).

MG-specific expression of TFs was achieved by a GFAP promoter(*22*) in the adeno-associated viruses (AAVs, serotype 9 or shH10) injected intravitreally or subretinally into the adult mouse eyes (Fig. 1B). Two weeks after injection, MG infected with GFAP-GFP AAVs displayed typical Müller cell morphology; however, the great majority of MG infected with GFAP-Math5-Brn3b-GFP AAVs changed their morphology, lost MG processes spanning the inner and outer retinal layers and migrated into the ganglion cell layer (GCL) (Fig. 1C). They were immunoreactive for RGC markers Rbpms or Brn3a and occasionally for the amacrine cell marker Tfap2a/2b, but not for the Müller cell marker Sox9 (Fig. 1C). Quantification of immunoreactive cells revealed that the Math5 and Brn3b combination reprogrammed infected MG into 92.8% Rbpms+ RGCs, 67.9% Brn3a+ RGCs, and 6.3% Tfap/2a/2b+ amacrine cells, whereas MG infected with control GFP AAVs remained as 100% Sox9+ Müller cells (Fig. 1G). Single Brn3b factor did not exert any reprogramming effect although single Math5 TF converted infected MG into 11.9% Rbpms+ RGCs, 6.6% Brn3a+ RGCs, and 14.8% Tfap/2a/2b+ amacrine cells (Fig. 1C, G). These results indicate that Math5 together with Brn3b are able to reprogram mature MG into RGCs with exceedingly high efficiency whereas either TF alone has no or limited capacity.

To further confirm MG reprogramming to RGCs by Math5 and Brn3b, we generated a Brn3b-GFP reporter mouse line in which GFP was simultaneously expressed with Brn3b to specifically mark most RGCs (Fig. 1D, F). Thus, most MG-derived RGCs in this line would be labeled by the GFP reporter. Indeed, within the GCL of retinas infected with GFAP-Math5-Brn3b-tdTomato AAVs, the majority of tdTomato-positive cells were also immunoreactive for GFP as well as Rbpms, whereas in control retinas, tdTomato-positive cells remained immunoreactive for Sox9 but negative for either GFP or Rbpms (Fig. 1E). Quantification of immunoreactive cells showed that 58.1% and 68.2% of tdTomato-positive cells were co-immunoreactive for GFP, 7 and 14 days post-infection with GFAP-Math5-Brn3b-tdTomato AAVs, respectively (Fig. 1H), confirming that RGCs were efficiently reprogrammed from MG by the action of both Math5 and Brn3b and that the reprogramming process was completed in a week in most cases. We next investigated whether MG proliferation was stimulated by Math5 and Brn3b before MG reprogramming took place. The retinas were labeled by EdU injection at day 2.5 or 4.5 following infection with GFAP-Math5-Brn3b-GFP AAVs, and harvested 1 or 4.5 days later (fig. S1A). In all cases, essentially no EdU-positive MG or other cells were observed (fig. S1), suggesting that Math5 together with Brn3b are able to reprogram mature MG into RGCs without triggering their proliferation.

Endogenous RGC axons project along a stereotypic visual pathway to connect to their appropriate central targets in the brain(*23*). We investigated whether MG-derived RGCs had the ability to form axons that follow the same pathway. At 3-4 weeks following infection of the adult retina by GFAP-Math5-Brn3b-tdTomato AAVs, numerous tdTomato-immunoreactive RGCs were observed on the vitreous surface of the retina, which extended axons that were fasciculated into many thick axon bundles immunoreactive for Tuj1 (Fig. 2A-E). These bundles extended all the way from the periphery through intermediate and central retinal areas to enter the optic disc (Fig. 2A-E). Once exiting the optic disc, these RGC axons, which were countless in number, continued to navigate through the optic nerve (Fig. 2F, G). The great majority of them crossed over the midline of the optic chiasm to continue their projection in the contralateral optic tract while a small number continued their projection in the ipsilateral optic tract (Fig. 2H-K). The predominant contralateral axon projection was confirmed by mono-ocular treatment with GFAP-Math5-Brn3b-tdTomato AAVs, which unambiguously showed that the great majority of axons from MG-derived RGCs crossed the midline of the optic chiasm (Fig. 2L). In the brain, there were plenty of tdTomato+ axons from MG-derived RGCs that reached and innervated various central targets responsible for both image-forming and non-image forming vision: the lateral geniculate nucleus, superior colliculus, olivary pretectal nucleus, terminal nucleus, accessory optic tract, and the above-mentioned optic chiasm (Fig. 2M-T). Therefore, MG-derived RGCs are able to extend long axons capable of navigating the entire visual pathway to innervate proper central targets.

**Fig. 2.**
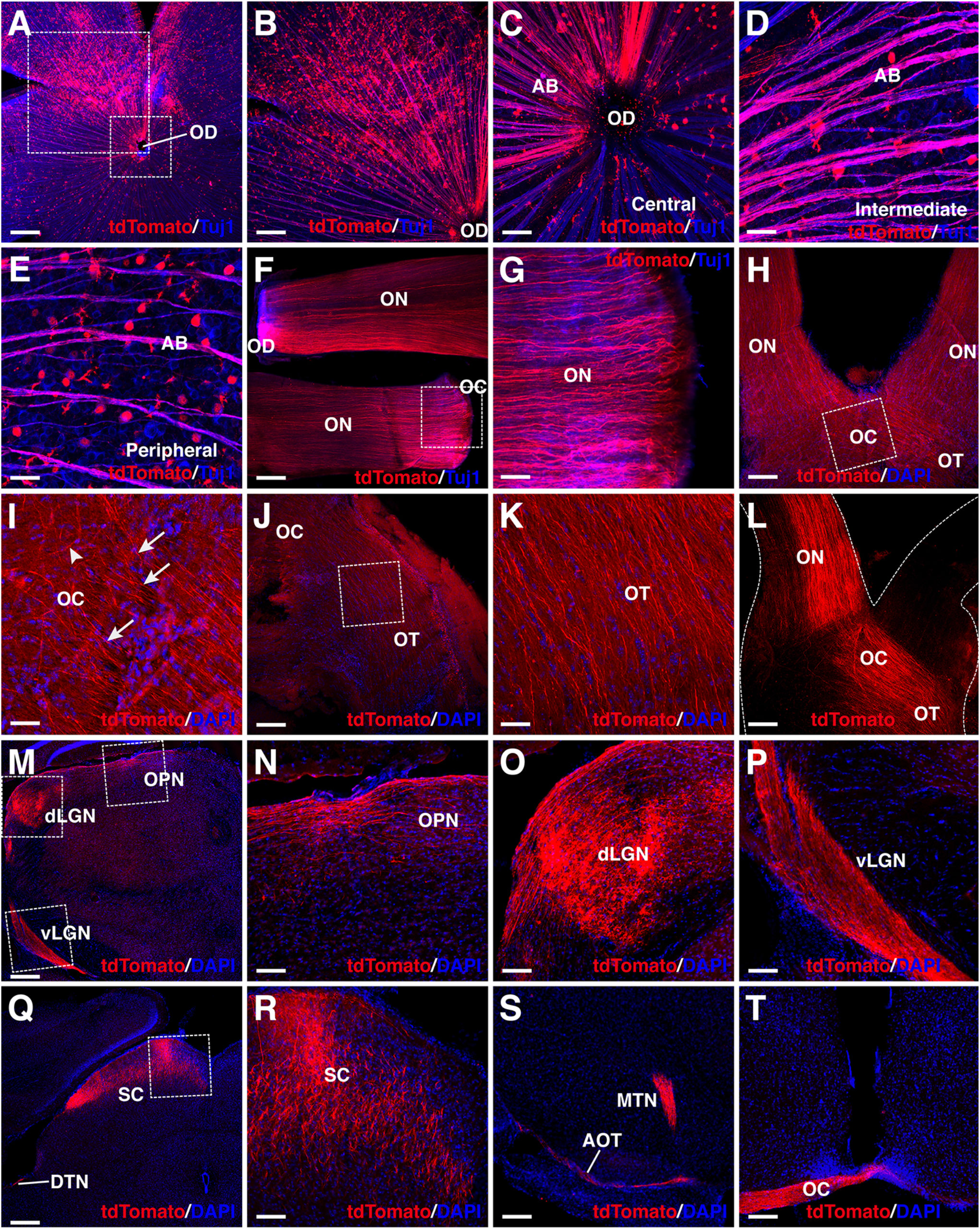
MG-derived RGCs recapitulate the visual projection pathway of endogenous RGCs. **(A-E**) Flat-mounts of adult mouse retinas treated with GFAP-Math5-Brn3b-tdTomato AAVs were double-immunolabeled with anti-tdTomato and anti-Tuj1 antibodies. The two areas outlined by the large and small squares in (A) are shown at a higher magnification in (B) and (C), respectively. Shown in (C-E) are representative images from the central, intermediate and peripheral retinas, respectively. (**F-K**) The optic nerves, optic chiasms and optic tracts of mice treated with GFAP-Math5-Brn3b-tdTomato AAVs were immunolabeled with an anti-tdTomato antibody and an anti-Tuj1 antibody (F,G) or counterstained with nuclear DAPI (H-K). Panel (F) shows one optic nerve segment near the optic disc (OD) and another near the optic chiasm (OC). Panels (G), (I) and (K) are higher magnification views of regions outlined in (F), (H) and (J), respectively. Arrows in (I) point to axons crossing the midline of the optic chiasm (contralateral projection) while the arrowhead indicates a non-crossing axon (ipsilateral projection). (**L**) Mono-ocular treatment of adult mice with GFAP-Math5-Brn3b-tdTomato AAVs revealed that tdTomato-immunoreactive axons projected predominantly into the contralateral optic tract. (**M-T)** Brain areas innervated by MG-derived RGCs in mice treated with GFAP-Math5-Brn3b-tdTomato AAVs. Shown in (N-P) are higher magnification views of the corresponding outlined regions in (M) and shown in (R) is a higher magnification view of the region outlined in (Q). Abbreviations: AB, axon bundle; AOT, accessory optic tract; dLGN, dorsal lateral geniculate nucleus; DTN, dorsal terminal nucleus; EF, MG endfoot; MTN, medial terminal nucleus; OC, optic chiasm; OD, optic disc; ON, optic nerve; OPN, olivary pretectal nucleus; OT, optic tract; RGC, retinal ganglion cell; SC, superior colliculus; vLGN, ventral lateral geniculate nucleus. Scale bars: 320 μm (A,M,Q), 160 μm (B, F, H, J, L, S, T), 80 μm (N, O, P, R), 40 μm (C-E, G, I, K).

To evaluate the functionality of regenerated RGCs, we attempted to reprogram MG into RGCs in adult *Brn3b^AP/AP^* knockout mutant mice where 70-80% of RGCs are lost(*17*, *24*) (Fig. 3). On the vitreous surface of mutant retinas infected with GFAP-tdTomato AAVs, except for numerous tdTomato-positive MG endfeet, there were no RGCs and axon bundles labeled by tdTomato (Fig. 3B, C). By contrast, in mutant retinas infected with GFAP-Math5-Brn3b-tdTomato AAVs, many tdTomato-immunoreactive RGCs were present and these regenerated RGCs extended numerous tdTomato-positive axon bundles that exhibited proper projection to the optic disc (Fig. 3D-G). Moreover, these nerve fibers navigated all the way through the optic nerve, optic chiasm and optic tract whereas no tdTomato-positive axons were seen in the control optic nerve (Fig. 3H-K). Overall, there was two to three-fold increase of RGCs in the central, intermediate and peripheral regions in retinas infected with GFAP-Math5-Brn3b-tdTomato AAVs (Fig. 3L, M; fig. S2). Thus, similar to in wild-type retinas, RGCs can be efficiently reprogrammed from MG in *Brn3b^AP/AP^* null mutant retinas as well.

**Fig. 3.**
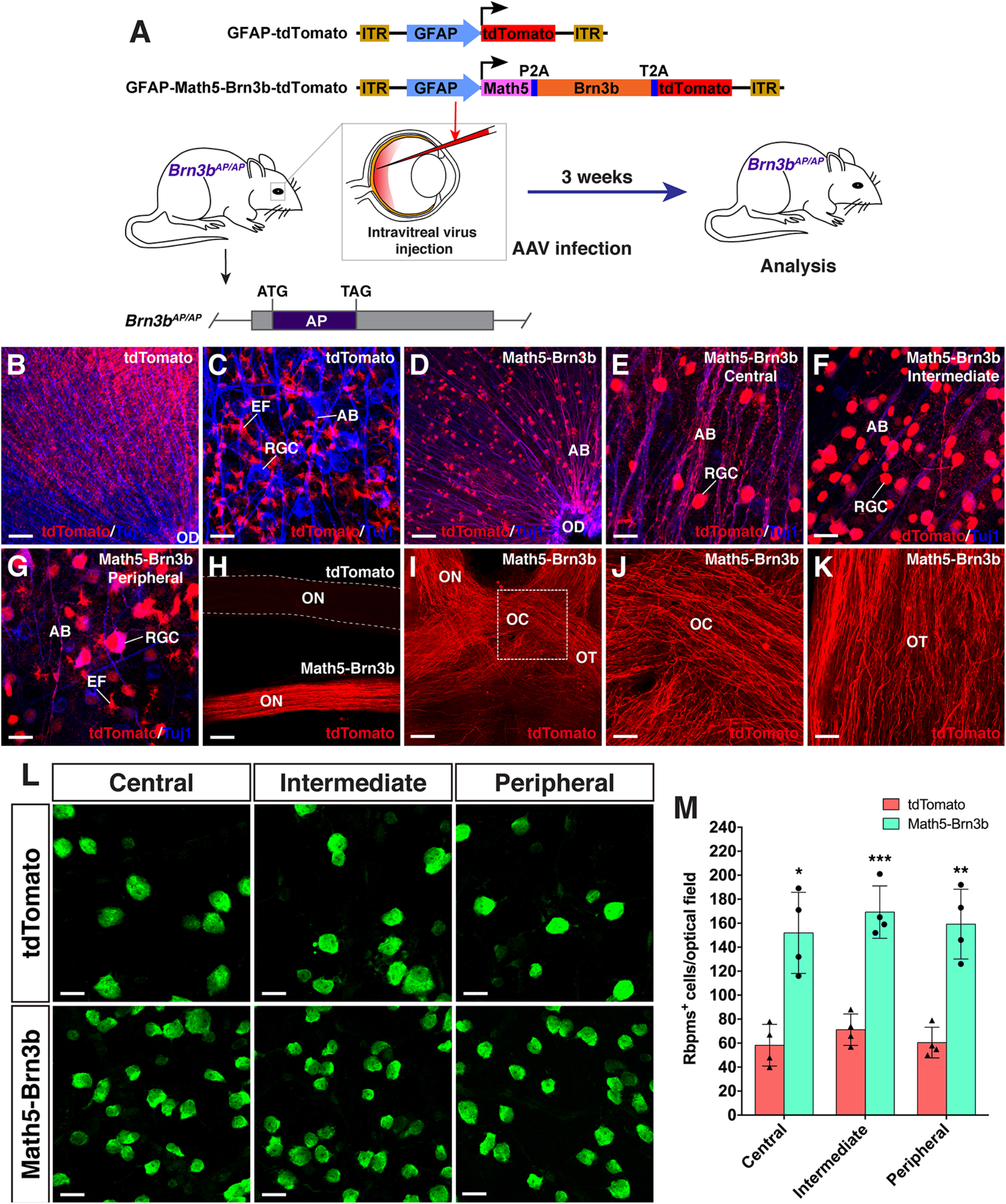
RGC regeneration in *Brn3b^AP/AP^* mice. (**A**) Schematic of the AAV constructs and infection procedure to regenerate RGCs in adult *Brn3b^AP/AP^* mice. (**B, C**) Flatmounts of *Brn3b^AP/AP^* retinas treated with GFAP-tdTomato AAVs were doubleimmunolabeled with anti-tdTomato and anti-Tuj1 antibodies. (**D-G**) Flat-mounts of *Brn3b^AP/AP^* retinas treated with GFAP-Math5-Brn3b-tdTomato AAVs were doubleimmunolabeled with anti-tdTomato and anti-Tuj1 antibodies. Shown in (E-G) are representative images from the central, intermediate and peripheral retinas, respectively. (**H**) The optic nerves from *Brn3b^AP/AP^* mice treated with GFAP-Math5-Brn3b-tdTomato AAVs were immunoreactive for tdTomato whereas those from *Brn3b^AP/AP^* mice treated with GFAP-tdTomato AAVs were not. (**I-K**) The optic nerves, optic chiasms and optic tracts from *Brn3b^AP/AP^* mice treated with GFAP-Math5-Brn3b-tdTomato AAVs were immunoreactive for tdTomato. (**L**) Flat-mounts of central, intermediate and peripheral *Brn3b^AP/AP^* retinas treated with GFAP-tdTomato or GFAP-Math5-Brn3b-tdTomato AAVs were immunostained with an anti-Rbpms antibody. (**M**) Quantification of Rbpms+ cells in central, intermediate and peripheral *Brn3b^AP/AP^* retinas treated with GFAP-tdTomato or GFAP-Math5-Brn3b-tdTomato AAVs. Data are presented as mean ± SD (n=4). Asterisks indicate significance in unpaired two-tailed Student’s t-test: *p<0.005, ***p<0.001, ***p<0.0005. Abbreviations: AB, axon bundle; EF, MG endfoot; OC, optic chiasm; OD, optic disc; ON, optic nerve; OT, optic tract; RGC, retinal ganglion cell. Scale bars: 114 μm (I), 80 μm (B, D, H), 40 μm (J, K), 20 μm (C, E-G, L).

To determine whether we reprogrammed MG into functional RGC neurons, we carried out whole-cell patch-clamp recording of RGCs in Brn3b-GFP reporter mouse retinas infected with GFAP-Math5-Brn3b-tdTomato AAVs, where most reprogrammed RGCs should be labeled by both tdTomato and GFP reporters. Indeed, we found cells that displayed both red and green fluorescence in the GCL (Fig. 4A), and that the great majority of them (10 out of 11) had action potential responses and exhibited excitatory postsynaptic potentials (Fig. 4B-D), suggesting that RGCs reprogrammed from MG are able to differentiate into mature functional neurons with characteristic physiological membrane properties. Given the apparent proper projection of the MG-derived RGCs along the visual pathway, we tested whether they were able to transmit light responses to the primary visual cortex in vivo. Visual evoked potentials (VEPs) to a flash light in the primary visual cortex were recorded from mouse models with the optic nerve crushed (ONC). From ONC eyes infected with GFAP-GFP AAVs (control), the light stimulus elicited much smaller VEP responses compared to those from normal wild-type eyes. Infection of the ONC eyes with GFAP-Math5-Brn3b-GFP AAVs (treated) triggered obviously stronger VEP responses than the control treatment (Fig. 4E). Quantification showed that the amplitudes of the VEP response peaks from the treated eyes were increased by ~ 67% compared to those from the control eyes (Fig. 4F). Additionally, we recorded VEPs from *Brn3b^AP/AP^* mice infected with GFAP-Math5-Brn3b-tdTomato AAVs (treated) or GFAP-tdTomato AAVs (control). In agreement with the loss of approximately 70% RGCs in *Brn3b* knockout animals(*17*), the control mutant eyes gave only small VEP responses (Fig. 4G). Again, the light stimulus triggered stronger VEP responses from the treated eyes, with the response amplitudes increased by ~ 44% compared to control eyes (Fig. 4H), consistent with the observed increase of RGCs in the treated retina (Fig. 3).

**Fig. 4.**
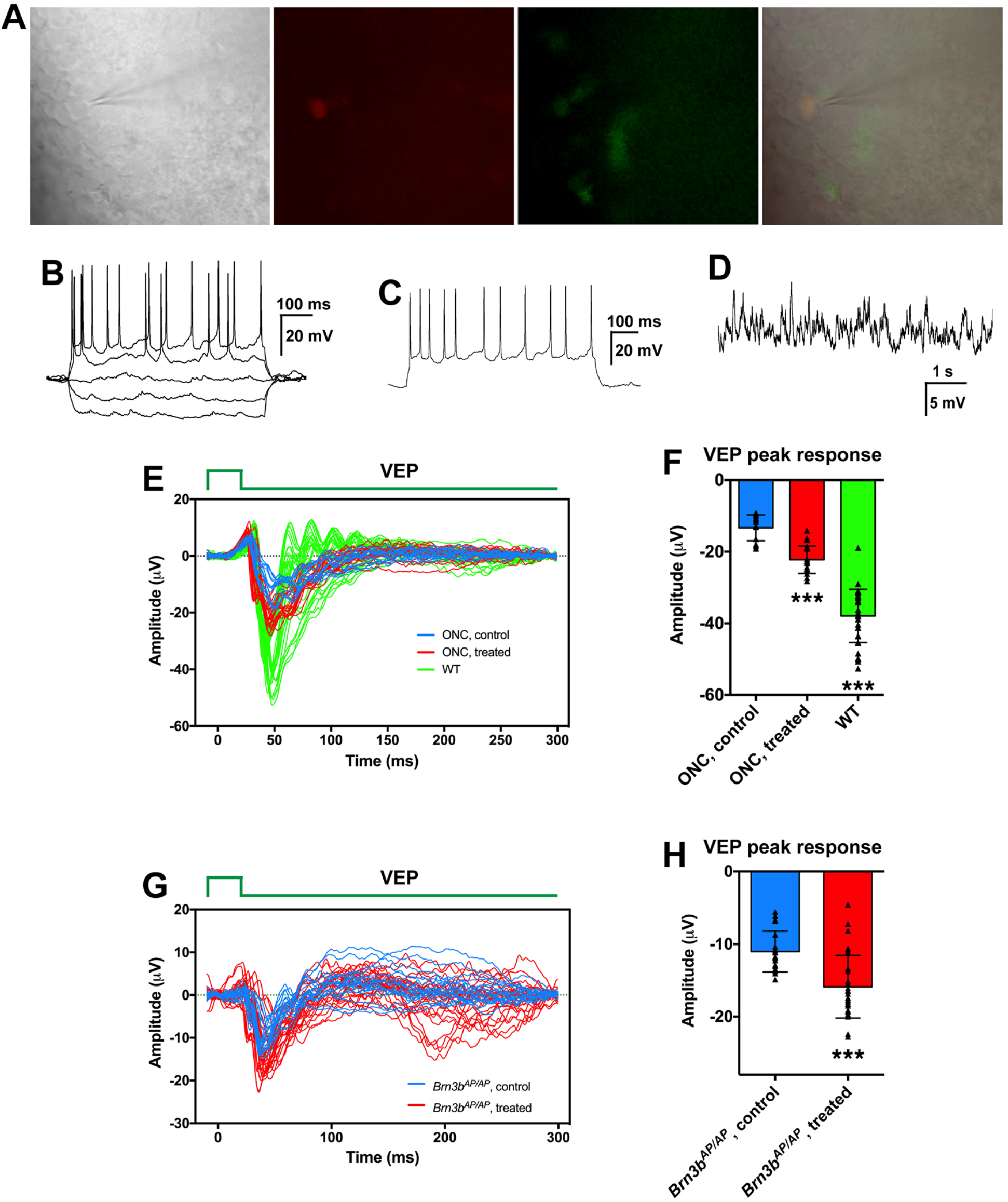
MG-derived RGCs improve visual function of glaucoma mouse models. **(A-D)** Current-clamp recording of RGCs in Brn3b-GFP reporter mouse retinas treated with GFAP-Math5-Brn3b-tdTomato AAVs. (A) An MG-derived RGC marked by both tdTomato (red) and GFP (green) was chosen for patch-clamp recording. (B) Current-clamp recordings revealed action potential responses of the MG-derived RGC under current injection. (C) Action potentials were induced after depolarization of the patched cell. (D) Excitatory postsynaptic potentials recorded from an MG-derived RGC. (**E, F**) Visual evoked responses (VEPs) to a flash light in the visual cortex of ONC mouse models in right eyes (WT, without optic nerve crush, n=7) and in left eyes (optic nerve crushed) treated with GFAP-Math5-Brn3b-GFP AAVs (ONC, treated, n=4) or GFAP-GFP AAVs (ONC, control, n=3). Shown in (E) are responses from all trials and five trials were performed for each eye. Shown in (**F)** are amplitudes of the VEP response peaks for control, treated and WT eye groups. Points represent single trials. Data are presented as mean ± SD. Asterisks indicate significance in unpaired two-tailed Student’s t-test: ***p<0.0001. (**G, H**) VEPs to a flash light in the visual cortex of *Brn3b^AP/AP^* mice treated with GFAP-Math5-Brn3b-tdTomato AAVs (*Brn3b^AP/AP^*, treated, n=6 eyes) or GFAP-tdTomato AAVs (*Brn3b^AP/AP^*, control, n=4 eyes). Shown in (G) are responses from all trials and five trials were performed for each eye. Shown in (**H)** are amplitudes of the positive VEP response peaks for control and treated eye groups. Points represent single trials. Data are presented as mean ± SD. Asterisks indicate significance in unpaired twotailed Student’s t-test: ***p<0.0001.

Given the demonstrated expression of all three Brn3 TF family members in RGCs and their functional redundancy(*25*–*29*), we investigated whether Brn3a and Brn3c have a similar reprogramming activity as Brn3b (figs. S3,S4). In adult wild-type retinas, Math5 combined with Brn3a or Brn3c converted many infected MG into Rbpms+ or Brn3b+ RGCs located within the GCL as well as some Tfap2a/2b+ amacrine cells located in the inner half of the inner nuclear layer and GCL (figs. S3B,S4B). Quantification revealed that Math5 combined with Brn3a reprogrammed MG into 77.6% Rbpms+ RGCs and 56.9% Brn3b+ RGCs and its combination with Brn3c converted MG into 51.7% Rbpms+ RGCs and 39.9% Brn3b+ RGCs (figs. S3C,S4C), whereas any of these TFs alone had no or much weaker reprogramming activity (figs. S3,S4). Thus, albeit weaker than Brn3b, Brn3a or Brn3c in combination with Math5 also exhibits a strong activity in reprogramming MG into RGCs.

Our data together demonstrate that mature MG, even without activation by injury or proliferation-stimulants, can be reprogrammed by developmentally-pertinent TFs to robustly generate functional RGCs. This is in contrast to previous observation of neurogenesis and rod generation by murine MG, which does require prior MG activation(*8*–*11*, *13*), suggesting a possible cell type-specificity of this requirement. The in vivo reprogrammed RGCs migrated into the GCL, made proper intra-retinal and extra-retinal projections through the entire visual pathway to form correct central connections, exhibited typical neuronal electrophysiological properties, and restored vision to glaucoma mouse models. These results implicate that even in the adult organism, the mammalian visual system may still maintain a relatively intact and permissive environment for regenerated RGC axons to outgrow and navigate to appropriate brain targets, and that unlike postnatal endogenous RGCs which lose their ability to extend(*30*), regenerated RGCs retain the ability to project axons along the visual pathway, similar to embryonically born RGCs(*30*). Given the many desirable feats of regenerated RGCs never achieved by transplanted retinal stem cells or in vitro differentiated RGCs, in vivo regeneration of RGCs by MG through directed reprogramming provides a promising new therapeutic approach to restore vision to patients with glaucoma and related optic neuropathies.

## Acknowledgements

We are grateful to Dr. William Klein (MD Anderson Cancer Center) for providing the *Brn3b^AP/+^* mice.

## Author contributions

D.X., K.J., S.L, Y.L. and M.X. conceived and designed the research. D.X., S.Q., X.H., R.Z., Q.L., W.H., H.C., B.G., and X.T. performed the experiments and analyzed the data. D.X., K.J., S.L. and M.X. interpreted the data and wrote the manuscript. All authors contributed to critical reading of the manuscript.

## Data and materials availability

All data from this study are available upon request.

