## Supplementary Materials for "Directed robust generation of functional retinal ganglion cells from Müller glia"

### Materials and Methods

**Animals.** All experiments on mice were performed according to the IACUC (Institutional Animal Care and Use Committee) standards, and approved by Sun Yat-sen University and Zhongshan Ophthalmic Center. The *Brn3b*$^{AP/AP}$ knockin mutant mice were previously generated and maintained in our laboratory(*1*). The Brn3b-GFP reporter mouse line was created using the CRISPR/Cas9 gene editing system and the related results will be published elsewhere. The C57BL6/J and CD1 mice were purchased from the Vital River Laboratories (Beijing, China). All genotypes were determined by PCR.

**Construction of viral plasmids and AAV production and injection.** A GFAP promoter(*2*) was subcloned into the pAAV-CAG-EGFP vector (Addgene plasmid # 37825) to replace CAG, generating the GFAP-GFP vector. The full-length ORFs of murine Math5, Brn3a and Brn3b alone or in combination via P2A or T2A were subsequently subcloned into the GFAP-GFP vector to construct desired AAV viral plasmids. By replacing GFP, we also used this vector as a backbone to construct the AAV plasmid expressing tdTomato.

AAV production was performed as described previously with modification(*3*). In brief, 20-24 hrs before transfection, AAV-293 cells (Sangon Biotech Co., Ltd, Shanghai, China) were cultured in 10 150-mm plates. For each plate, they were transfected with 6 μg of AAV vector DNA, 6 μg of AAV Rep/Cap plasmid DNA and 18 μg of adenovirus helper plasmid DNA using the Polyethylenimine (PEI) transfection method. After 60-72 hrs, the transfected cells were collected and resuspended in lysis buffer (150 mM NaCl, 20 mM Tris-HCl, pH8.0), then for three times frozen in dry ice/ethanol bath and thawed completely in 55°C water bath. The cell lysate was digested with Benzonase (50 U/ml) for 1 hr at 37°C, and centrifuged to remove the cell debris. For purification, the virus-containing supernatant was applied to discontinuous iodixanol gradients followed by ultracentrifugation. The virus band was collected from the 40%–60% interface using a syringe with a 21-gauge needle. The iodixanol solution was exchanged to 1xDPBS (Dulbecco's phosphate-buffered saline) using the Amicon Ultra-15 centrifugal filter units from Millipore and the viruses were further concentrated by shrinking the volume. Virus titers were determined by qRT-PCR using linearized plasmid standards and primers against the ITR. AAVs were injected intravitreally or subretinally into adult mouse eyes using a microsyringe with a 33-gauge needle as described(*4, 5*).

**Electrophysiological recording.** Three weeks following infection of the eyes of adult Brn3b-GFP reporter mice by GFAP-Math5-Brn3b-tdTomato AAVs, the retina was dissected out from the eyeball and its edge was removed to allow the tissue to lie flat. It was transferred to a recording chamber and bathed in external solution containing the Ames' medium (Sigma-Aldrich). The chamber was mounted on a microscope equipped with a 40× water immersion objective. The cells and recording pipettes were viewed on a monitor that was coupled to a camera (Scientifica SciCam Pro, Canada). Oxygenated external solution was continuously perfused into the recording chamber at a flow rate of 1.5–2 ml/min. GFP-positive and tdTomato-positive cells were identified with a mercury lamp (TH4-200, Olympus, Japan). Then, whole-cell patch-clamp recordings were performed with a 700B amplifier (Molecular Devices, USA) and digitized at 10 kHz with a Digidata 1550B (Molecular Devices, USA). The responses of the cells were recorded with 6–9 MΩ resistance pipettes that were filled with an internal solution consisting of the following: 123 mM K-gluconate, 12 mM KCl, 10 mM HEPES, 0.2 mM EGTA, 4 mM Mg-ATP, 0.3 mM Na-GTP,10 mM $Na_2$-phosphocreatine, and 20 μg/ml glycogen (the pH value was adjusted to 7.25 with 0.5 M KOH). For current-clamp recording, we set the initial resting membrane potential (V-rest) to -70 mV using a small, constant holding current and applied current pulses with a step size of 20 pA to test the ability to generate action potentials.

**Optic nerve crush (ONC) injury.** Mice were anesthetized by intraperitoneal injection of 4% chloral hydrate and one drop of 0.4% oxybuprocaine hydrochloride was administered for local anesthesia. ONC was performed as described(*6*). Briefly, a small incision was made with scissors in the conjunctiva of the left eye located at the 3-9 o'clock of eyeball. The exposed optic nerve was grasped approximately 1 mm from the eyeball with forceps for 10 s. It was then released to allow the eyeball to rotate back into place. Three days following ONC, the left eye of each animal was infected with AAVs by intravitreal injection.

**Visual evoked potential (VEP) test.** Four to six weeks after AAV injection, the ONC and

*Brn3b*$^{AP/AP}$ mice were dark-adapted overnight, prior to being prepared for the experiments. They were anesthetized by intraperitoneal injection of 4% chloral hydrate and their pupils were dilated with a drop of tropicamide. One drop of 0.4% oxybuprocaine hydrochloride was administered for local anesthesia of the cornea.

During VEP recordings which were carried out using the Celeris ERG system (Diagnosys LLC, MA, USA), the animals were placed on a heated platform to keep warm. Needle electrodes placed subcutaneously at the base of the tail and at the snout served as ground and reference electrodes, respectively. The active electrode was inserted subcutaneously at the midline at the back of the head. Two contact-lens light-emitting diodes (LEDs) were placed over the two eyes of the animal to serve as light stimulators. Each eye was separately exposed to white light flashes of 0.05 cd.s/m$^2$, swept 100 times per trial. For each mouse, we performed five trials. Analyses were performed using GraphPad Prism 7.

**Immunohistochemistry.** Immunostaining of retinal sections, retinal flat-mounts and optic nerves were carried out as previously described(*7, 8*). Mouse brain tissues were immunostained as free floating sections also as previously described(*9*). In brief, for section labeling, retinas were fixed in 4% paraformaldehyde (PFA) in PBS for 30 min at 4 °C and sectioned at 14 μm. Sample sections were washed 3 times with 0.1% Tween in PBS (PBST) for 5 min each before being incubated in 5% normal donkey serum in PBST for 1 h at room temperature (RT). Then primary antibodies in 2% normal donkey serum in PBST were added for overnight incubation at 4 °C. After washing with PBST, the sections were incubated with secondary antibodies and DAPI in PBST for 1 hr at RT. Images were captured by a laser scanning confocal microscope (Carl Zeiss, LSM700).

The following primary antibodies were used: GFP (abcam, ab6673, 1:2000), RFP (Rockland antibodies & assays, 40657, 1:1000), RBPMS (Novus, NBP2-20112, 1:1000), Brn3a (Millipore, MAB1585, 1:500), Brn3b (Santa Cruz, SC-390780, 1:1000), Sox9 (Millipore, ab5535, 1:1000), Tfap2a&b (abcam, ab11828, 1:1000), and Tuj1 (Covance, MMS-435P, 1:1000).

**EdU staining.** At 2.5 days and 4.5 days following infection of adult mouse retinas with GFAP-

Math5-Brn3b-GFP AAVs, 2 μl of EdU solution (1 mg/ml) was injected into the vitreous chamber of each eye. One day or 4.5 days later, under deep anesthesia induced with intraperitoneal injection of chloral hydrate (4.5 μg/g body weight), mice were intracardially perfused with cold PBS for 5 min, followed by 4% cold PFA in PBS for 15 min. The eyeballs were isolated and post-fixed in 4% PFA for 1 hr at 4 °C. The retinas were dissected out and the vitreous was completely removed. They were shaped into a "petal" by 4 to 5 radial incisions, flattened in a 48-well plate, and permeabilized with 0.3% Triton-100 in PBST for 15 min at RT. After incubation in 10% normal donkey serum in PBST for 2 hrs at RT, the retinas were incubated in primary antibodies against Sox9 and GFP diluted in 2% normal donkey serum in PBST for 2 days at 4 °C. Retinas were washed with PBST and incubated with secondary antibodies for 2 hrs. After three washes with PBS, the retinas were stained for EdU with the Click-iT EdU Kit (Invitrogen). Images were captured by a laser scanning confocal microscope (Carl Zeiss, LSM700).

**Statistics.** Statistical analysis was performed using the GraphPad Prism 7 and Microsoft Excel computer programs. The results are expressed as mean ± SD for experiments conducted at least in triplicates. Unpaired two-tailed Student's t-test and/or Mann-Whitney test was used to assess differences between two groups, and a value of $P < 0.05$ was considered statistically significant.

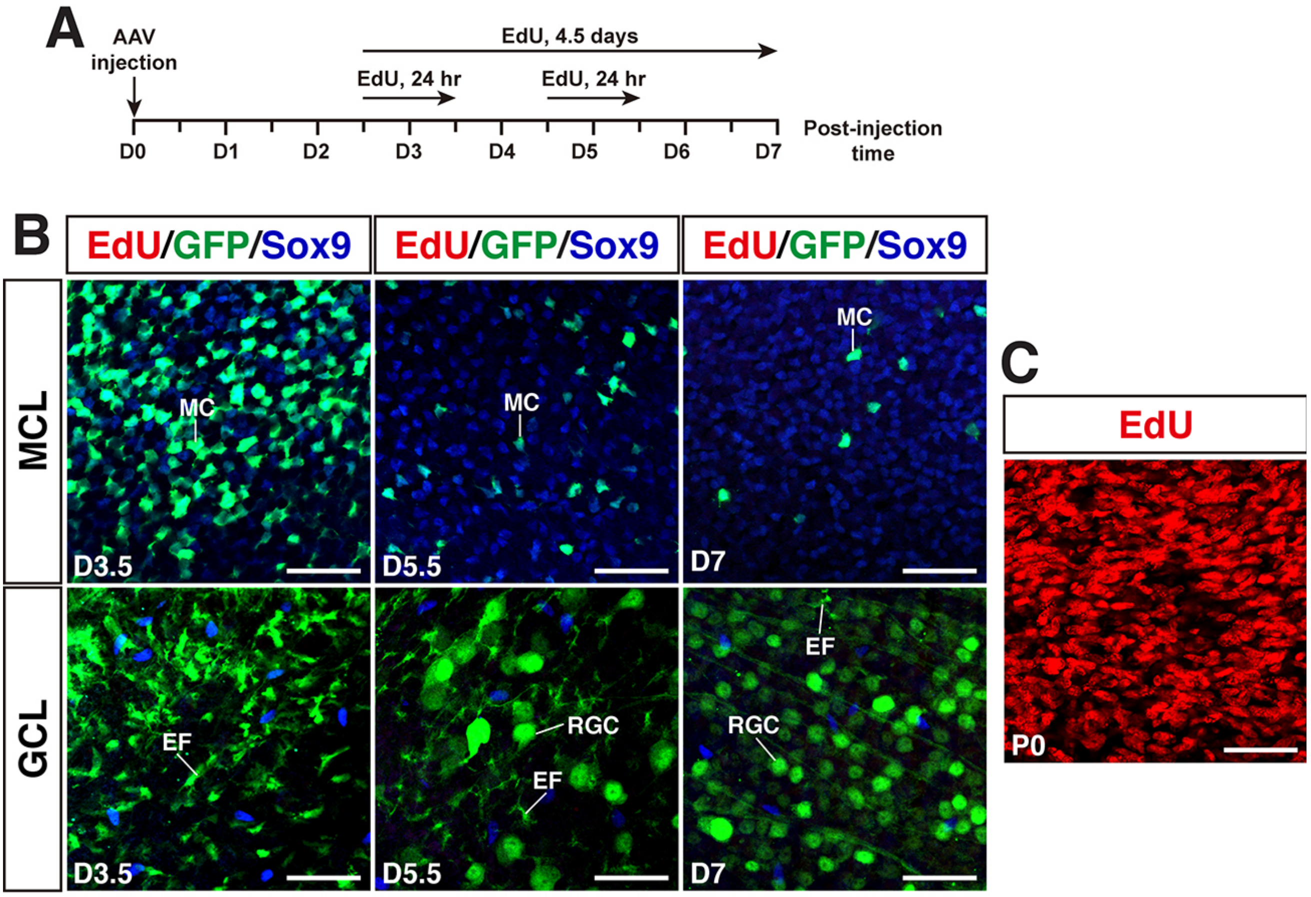


**Fig. S1. Math5 and Brn3b together do not trigger proliferation of mature MG.** (**A**) Schematic of EdU labeling schedule following infection of adult mouse retinas with GFAP-Math5-Brn3b-GFP AAVs. (**B**) Flat-mounts of adult mouse retinas infected with AAVs and pulse-labeled by EdU were fluorescently stained for EdU and immunostained for both GFP and Sox9. The confocal images are focused on the Müller cell layer (MCL) or ganglion cell layer (GCL). There are no EdU-positive cells present. (**C**) Flat-mounts of P0 mouse retinas pulse-labeled by EdU were fluorescently stained for EdU. There are numerous EdU-positive cells present. Abbreviations: EF, MG endfoot; MC, Müller cell; RGC, retinal ganglion cell. Scale bars: 40 $\mu$m (B,C).

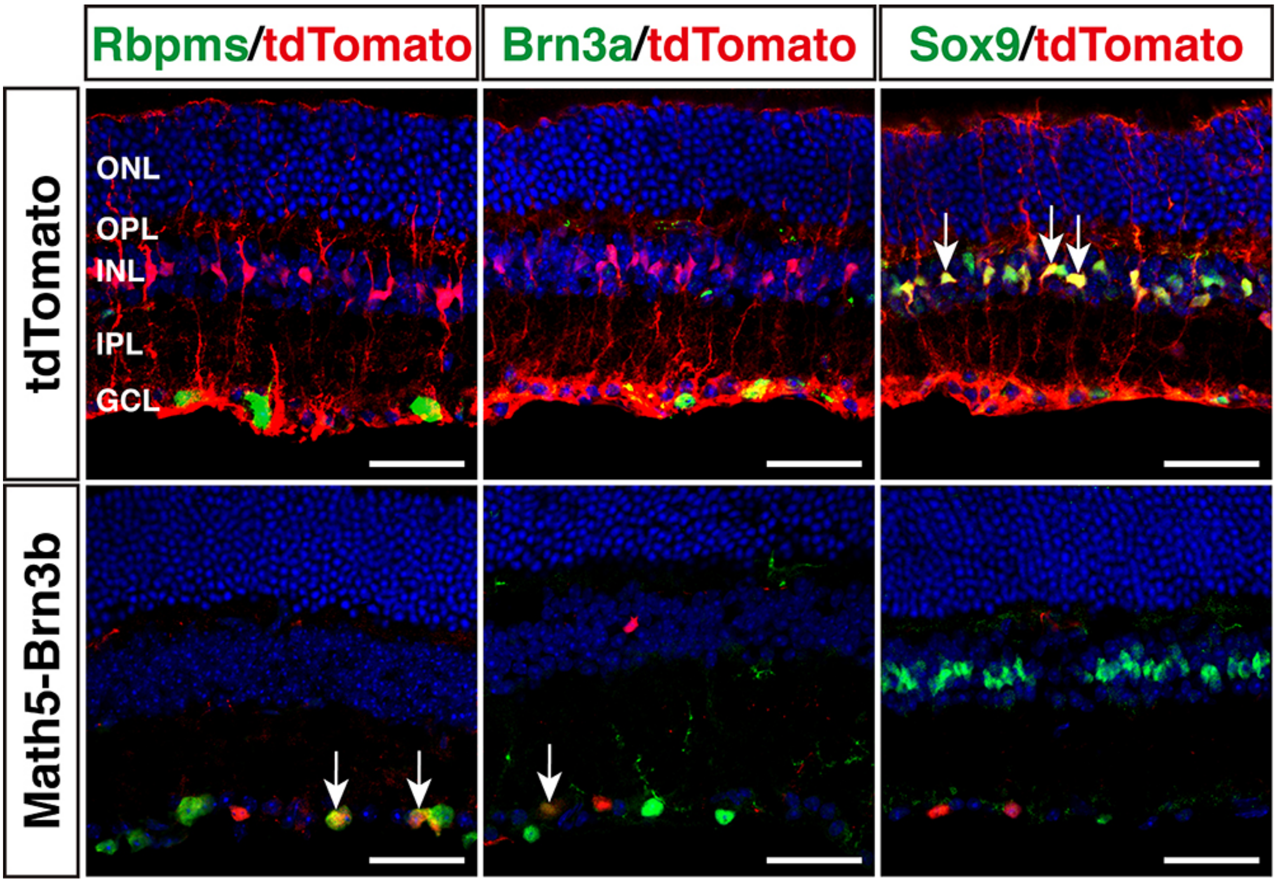


**Fig. S2. RGC regeneration in the *Brn3b$^{AP/AP}$* retina.** Sections from *Brn3b$^{AP/AP}$* retinas infected with GFAP-tdTomato or GFAP-Math5-Brn3b-tdTomato AAVs were double-immunostained with antibodies against tdTomato and Rbpms, Brn3a or Sox9. They were also counterstained with nuclear DAPI. Arrows point to representative colabeled cells. Compared to the control, Math5 and Brn3b together increased RGCs co-immunoreactive for both tdTomato and Rbpms or Brn3a. Abbreviations: GCL, ganglion cell layer; INL, inner nuclear layer; IPL, inner plexiform layer; ONL, outer nuclear layer; OPL, outer plexiform layer. Scale bars: 40 μm.

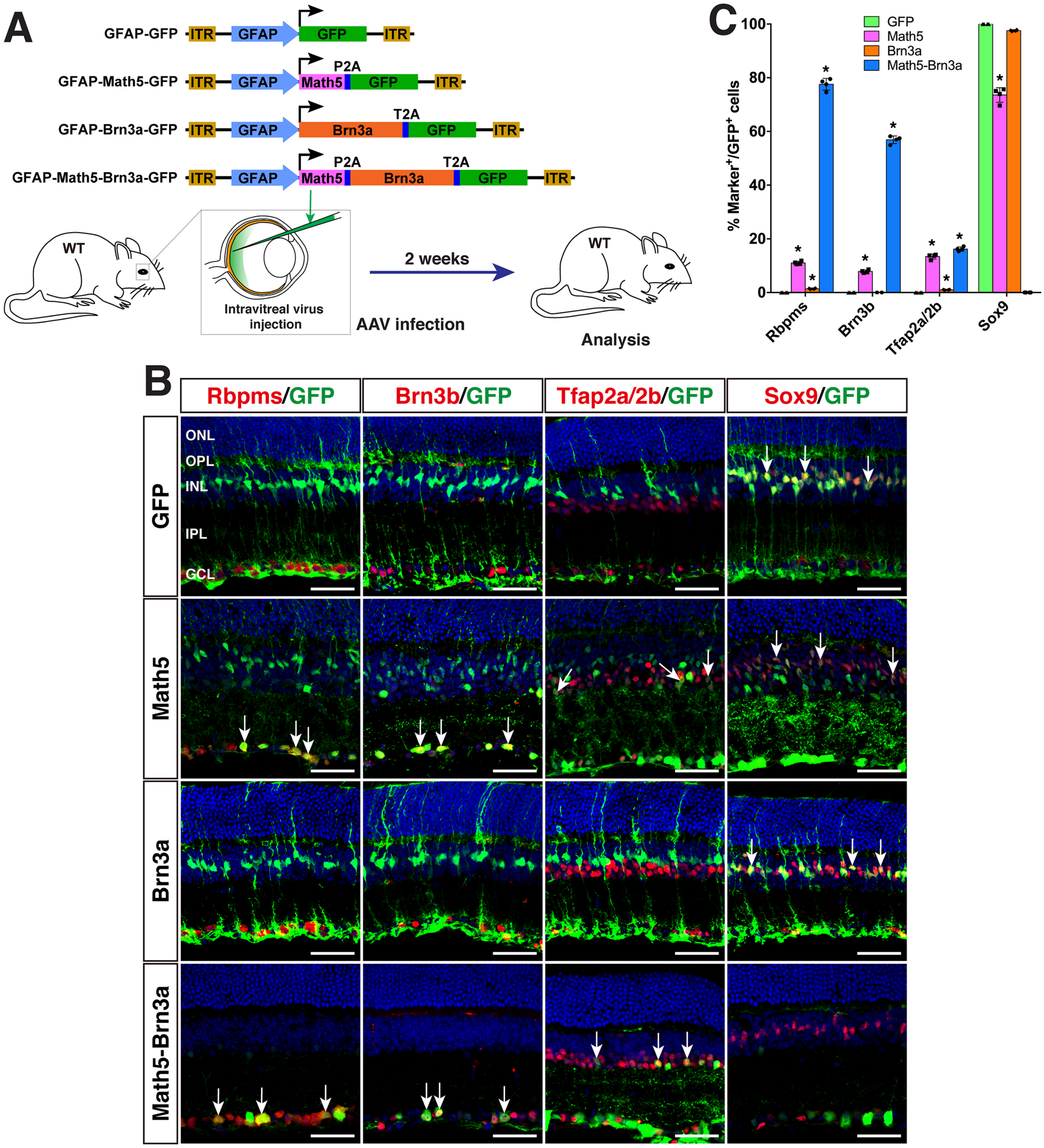


**Fig. S3. Generation of RGCs by reprogramming adult mouse MG with Math5 and Brn3a.** (**A**) Schematic of the AAV constructs and infection procedure to generate RGCs in wild-type (WT) mice. (**B**) Two weeks after intravitreal injection of GFAP-GFP, GFAP-Math5-GFP, GFAP-Brn3a-GFP, or GFAP-Math5-Brn3a-GFP AAVs, sections from infected retinas were double-immunolabeled with the indicated antibodies and counterstained with nuclear DAPI. Arrows point to representative colabeled cells. (**C**) Quantitation of GFP+ cells that become immunoreactive for Rbpms, Brn3b, Tfap2a/2b or Sox9 in retinas infected with GFAP-GFP, GFAP-Math5-GFP, GFAP-Brn3a-GFP, or GFAP-Math5-Brn3a-GFP AAVs. Data are presented as mean ± SD (n=4).

Asterisks indicate significance in unpaired two-tailed Student′s t-test: *$p<0.0001$. Abbreviations: GCL, ganglion cell layer; INL, inner nuclear layer; IPL, inner plexiform layer; ONL, outer nuclear layer; OPL, outer plexiform layer. Scale bars: 40 μm (B).

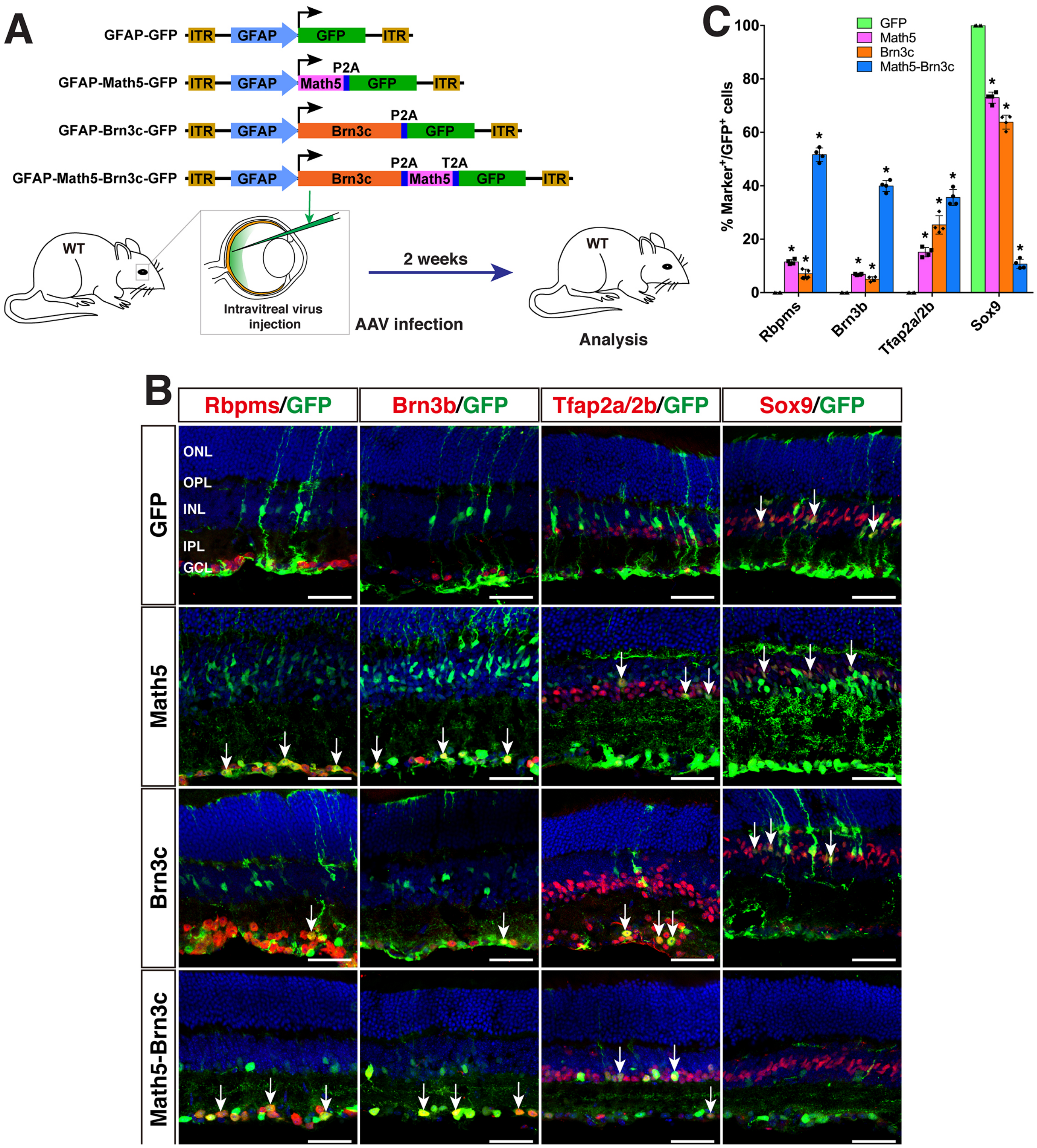


**Fig. S4. Generation of RGCs by reprogramming adult mouse MG with Math5 and Brn3c.** (**A**) Schematic of the AAV constructs and infection procedure to generate RGCs in wild-type (WT) mice. (**B**) Two weeks after intravitreal injection of GFAP-GFP, GFAP-Math5-GFP, GFAP-Brn3c-GFP, or GFAP-Math5-Brn3c-GFP AAVs, sections from infected retinas were double-immunolabeled with the indicated antibodies and counterstained with nuclear DAPI. Arrows point to representative colabeled cells. (**C**) Quantitation of GFP+ cells that become immunoreactive for Rbpms, Brn3b, Tfap2a/2b or Sox9 in retinas infected with GFAP-GFP, GFAP-Math5-GFP, GFAP-Brn3c-GFP, or GFAP-Math5-Brn3c-GFP AAVs. Data are presented as mean ± SD (n=4).

Asterisks indicate significance in unpaired two-tailed Student's t-test: *$p<0.0005$. Abbreviations: GCL, ganglion cell layer; INL, inner nuclear layer; IPL, inner plexiform layer; ONL, outer nuclear layer; OPL, outer plexiform layer. Scale bars: 40 μm (B).
